# SANe: The Seed Active Network for Discovering Transcriptional Regulatory Programs of Seed Development

**DOI:** 10.1101/165894

**Authors:** Chirag Gupta, Arjun Krishnan, Andrew Schneider, Cynthia Denbow, Eva Collakova, Pawel Wolinski, Andy Pereira

## Abstract

Developing seeds undergo coordinated physiological and morphological changes crucial for development of the embryo, dormancy and germination. The metabolic changes that occur during seed development are regulated by interconnected network of Transcription Factors (TFs) that regulate gene expression in a spatiotemporal manner. The complexity of these networks is such that the TFs that play key regulatory roles during seed development are largely unknown. In this study, we created a genome-scale regulatory network dedicated to describing regulation of biological processes within various compartments and developmental stages of Arabidopsis seeds. Differential network analysis revealed key TFs that rewire their targeting patterns specifically during seed development, many of which were already known, and a few novel ones that we verified experimentally. Our method shows that a high-resolution tissue-specific transcriptome dataset can be accurately modeled as a functional regulatory network predictive of related TFs. We provide an easy to use webtool using which researchers can upload a newly generated transcriptome and identify key TFs important to their dataset as well as gauge their regulatory effect on phenotypes observed in the experiment. We refer to this network as Seed Active Network (SANe) and made it accessible at https://plantstress-pereira.uark.edu/SANe/. We anticipate SANe will facilitate the discovery of TFs yet unknown for their involvement in seed related metabolic pathways and provide an interface to generate new hypothesis for experimentation.

## Introduction

The evolutionary success of plants lies in their ability to produce seeds and their dispersal which facilitates the progression of propagation. Seeds are complex structures that help plants halt their life cycle under unfavorable conditions and resume growth once the environmental conditions become favorable. In Arabidopsis, a double fertilization event marks the beginning of seed development that progresses into the development of embryo, endosperm and seed coat over a period of 20-21 days after pollination. These morphologically distinct sub-compartments within a seed play diverse roles and function in concert during the entire phase of seed formation. During maturation, synthesis of storage reserves occurs and traits such as desiccation tolerance and dormancy are acquired. These seed storage reserves fuel for seedling emergence during germination. Identifying genes that play important roles in seed development, especially the ones that are involved in carbon metabolism during the synthesis of storage reserves will greatly benefit the development of plant varieties with enhanced yield traits in the future.

Delineating regulatory networks is critical to identifying genes that regulate metabolic processes and underlying seed development and maturation. Several transcription factors (TFs) that regulate various aspects of seed development as well as germination have been revealed by genetic screens (Grossniklaus et al., 1998; Lotan et al., 1998; Ogas et al., 1999; Johnson et al., 2002; To et al., 2006). Among these TFs, three members of the B3 super family namely, *LEAFY COTYLEDON 2 (LEC2), ABSCISIC ACID INDENSITIVE 3 (ABI3)* and *FUSCA3 (FUS3)*, along with two members of the LEC1-type, *LEC1* and *LEC1-LIKE*, that together form the ‘LAFL’ network (Jia et al., 2013), are the most prominent players of seed maturation. However, the existing LAFL network is still incomplete and represents only a subset of regulatory networks active during seed development. The functional roles of several other seed-expressed TFs remain largely unknown. Although the genetic interactions, functional redundancy and cooperativity between TFs will be more accurately revealed by genetic interventions, delineating seed gene regulatory networks from a computational standpoint is critical to narrowing the search space for identification and prioritization of candidates for experimentation *in vivo*.

Expression-based network discovery from integrated transcriptome datasets is still the most widely used measure to gauge the association of uncharacterized genes to functional categories consisting of known genes (Obayashi and Kinoshita, 2011; Sato et al., 2012; Yim et al., 2013; Aoki et al., 2016; Krishnan et al., 2017). However, a typical coexpression network, by design, does not explicitly reveal information about the roles of regulatory genes in the observed expression patterns. Regulatory genes, for example genes encoding TFs, have been shown to regulate the expression of multiple target genes with common functions in the cell (Yu et al., 2011; Ambavaram et al., 2014). Because TFs themselves are transcriptionally regulated, they can also be targets of other TFs, giving the network a hierarchical organization (Ma et al., 2004; Spitz and Furlong, 2012), which makes it difficult to distinguish between direct and indirect regulatory edges as the signal propagates through the network. Moreover, the affinity of a TF for a target gene can be dependent on cell type and/or on the metabolic needs of the cell (Li et al., 2015). Owing to this complexity of transcriptional regulatory networks, inferring TF function using coexpression data can yield a high number of spurious interactions and should be monitored carefully. Accuracy in function prediction tasks can be improved by setting a unifying biological context on the underlying expression data (e.g., datasets for a specific tissue or condition), and filtering potential indirect regulatory edges using sophisticated reverse engineering algorithms (Basso et al., 2005; Faith et al., 2007; Huynh-Thu et al., 2010). A few previous studies have used these approaches to mine local (Fu and Xue, 2010; Ambavaram et al., 2014; Ramegowda et al., 2015; Ramegowda et al., 2017) and global context-specific regulatory networks in plants (Vermeirssen et al., 2014; Wilkins et al., 2016).

Here we used a seed-specific expression dataset to infer regulatory networks that manifest during seed development in Arabidopsis. Our network analysis method first defines functionally coherent modules (gene sets) with unique spatial and temporal expression patterns during different stages of embryo development. Then we used a combination of reverse-engineering and module enrichment analysis to find TFs that most likely govern the functional programs presented by these gene modules. We do not rely on the prediction of one-on-one TF-target interaction estimates, which typically requires selection of a not-easily-identifiable threshold score above which predicted TF-target interactions are deemed ‘true’. Instead, our method uses a weighted-analysis framework where predictions are made directly for the functional programs a TF is likely to regulate in the biological context under study, essentially exhausting the information content in the underlying tissue-specific dataset (Fig. 1).

**Figure 1:**
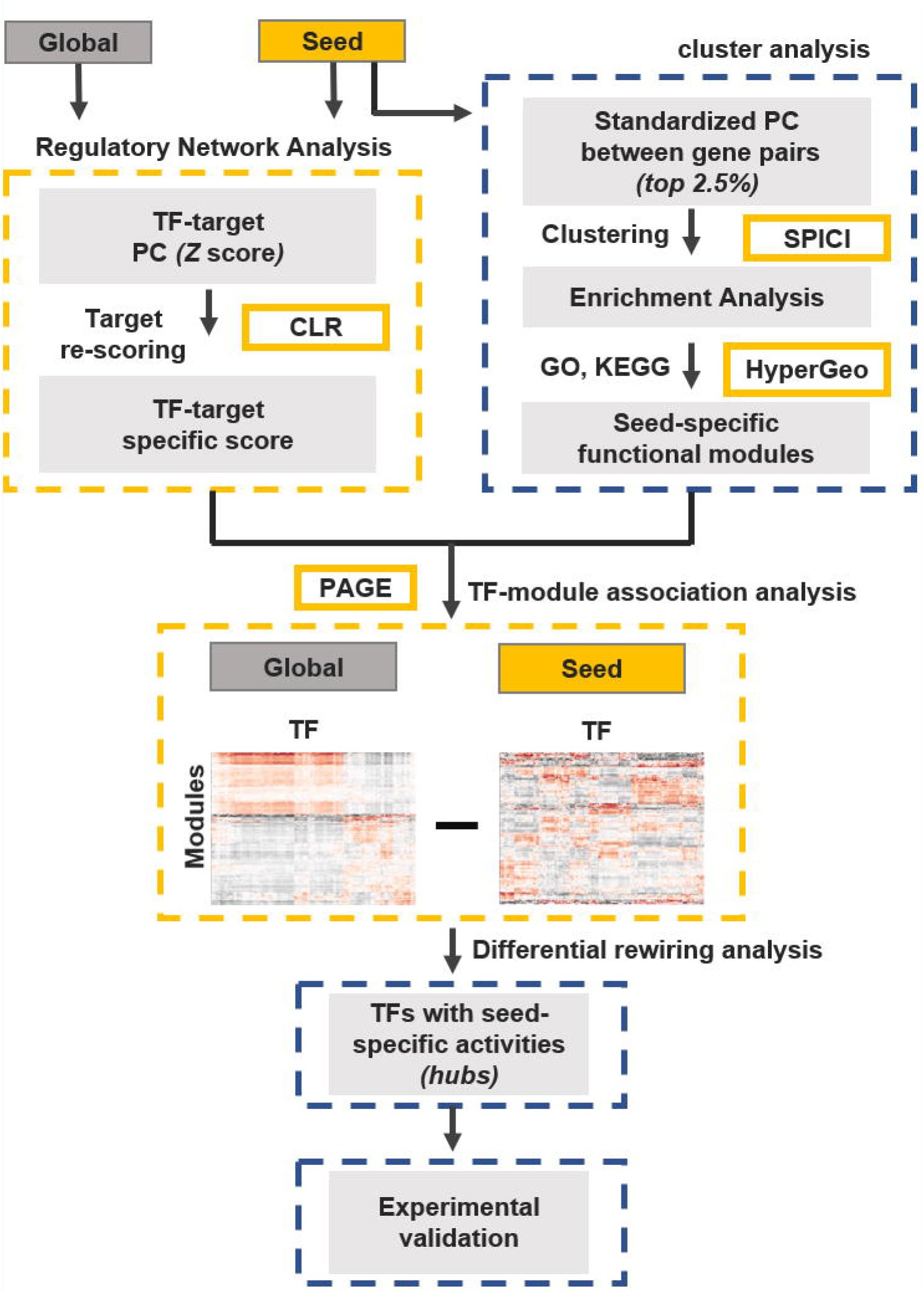
Pipeline for tissue-specific module regulatory network analysis. Two separate Arabidopsis gene expression compendiums (EC) were created: one from a seed-specific expression data series (GSE12404) and another from non-tissue-specific (global) expression datasets comprising of 140 samples. Datasets in both EC were normalized individually using RMA algorithm in R. Z scores of Pearson’s Correlations were calculated for all gene-pairs in both EC. From seed EC (top-right box with blue dashed lines), top 2.5% coexpressed gene pairs were connected to create a seed coexpression network which was clustered using SPICi at a range of clustering thresholds (*T_d_*), and an optimal clustering parameter was chosen based on genome coverage and coherence of genes as a functional group. 1563 clusters obtained at *T_d_* 0.80 were tested for enrichment of biological processes using gene ontology and known plant *cis* regulatory elements from multiple databases. A list of 1921 TFs was supplied to the CLR (context likelihood of relatedness) algorithm to predict their targets in both the EC separately (top-left box with orange dashed lines). The CLR-weighted targets of TFs in both networks were then subjected to enrichment analysis using seed modules as gene sets, and two TF-module association networks were created, one for the seed data and the other for the global data (middle box with orange dashed lines). The resulting module-regulatory networks were compared to each other and TFs with significant differences in their association scores with seed modules between the two networks were identified and subjected to experimental validations (bottom boxes with blue dashed lines). The specific algorithm/software used for analysis are indicated inside solid orange rectangles. PC: Pearson’s Correlation; SPICi: Speed and Performance In Clustering; CLR: Context Likelihood of Relatedness; HyperGeo: Hypergeometric test for testing the significance of overlap between two given gene sets; PAGE: Parametric Analysis of Geneset Enrichment.

Our method accurately recovered known functional programs of seed TFs reported in the literature for most of the cases tested. For example, a recently discovered association between the TF *AGL67* and desiccation tolerance (González-Morales et al., 2016), and *MYB107* and suberin (Lashbrooke et al., 2016) was correctly predicted in our network. These and several other correctly predicted associations (described later in the text) motivated us to create an online resource hosted at https://plantstress-pereira.uark.edu/SANe/ and integrated with tools to search the network and generate hypothesis for targeted downstream experiments.

## Results

### Seed coexpression network

To avoid implementing procedures of minimizing batch effects, genotype-specific variations and other errors associated with microarray data integration (Chen et al., 2011; Nygaard et al., 2015), we used the seed expression data derived in a single experiment (GSE12404) for our analysis. This series is comprised of a total of 87 samples derived from 6 discrete stages of seed development, and 5-6 different compartments within each stage, reflecting the most comprehensive source of seed-specific gene expression profiles (Belmonte et al., 2013).

Normalized coexpression scores between all gene pairs in the dataset were estimated and a coexpression network was created by connecting genes (nodes) where each edge represented statistically significant coexpression in the seed (see Methods). Top coexpressed edges were then used to cluster genes using an edge-weighted density-based graph clustering algorithm to identify communities, or groups of functionally similar genes in the network (Jiang and Singh, 2010). Rather than an arbitrary selection of the density threshold (*T_d_*) required by the clustering algorithm (to estimate ‘clique-ness’ of clusters), we sought to identify an optimum density threshold that yields clusters at a granularity that delivers biological information while preserving the inherent topological features of the network (Krishnan et al., 2017). A range of *T_d_* values were tested for performance in loss or gain of information, which was evaluated based on four criteria: 1) the trade-off between the number of clusters obtained at each *T_d_* and the fraction of original genes retained in those clusters, 2) The average modularity between clusters indicating how well genes within each cluster interact with each other as compared to genes outside their respective cluster (Albert, 2005), 3) the total number of Gene Ontology (GO) Biology Process (BP) terms represented (significantly enriched) by genes within all clusters, 4) and the average enrichment scores of all enriched BPs to detect if they were just ‘border-line’ significant or the overlap was large. Clearly, a *T_d_* value of 0.80 presented itself as the best parameter: 84% of all genes on the microarray formed the maximum of 1563 clusters (Supplemental Data S1), after which a significant loss of information occurred, as indicated by a sharp fall in the fraction of total genes retained (Fig. 2A). At the same threshold of 0.80, the average modularity within clusters was also maximized (at a bearable cost of gene loss) (Fig. 2B). The functional coherence of the network was also best preserved at *T_d_* 0.80 (Fig. 2C and 2D).

**Figure 2:**
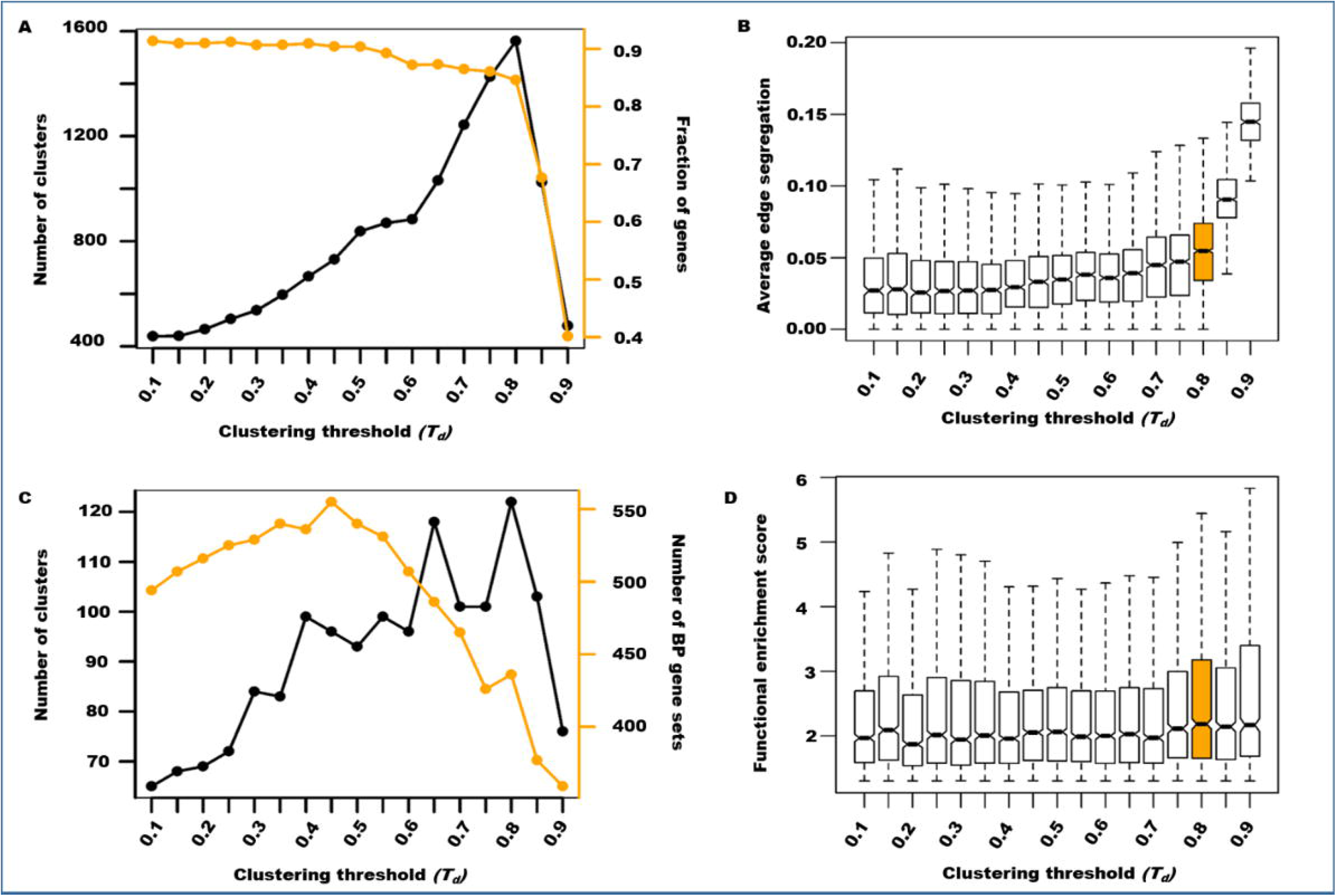
Evaluation of clustering threshold (*T_d_*). Genes from the Arabidopsis seed coexpression network were clustered at a range of *T_d_* values shown on the X axis in all figures. Each *T_d_* was examined by: A) A genome coverage plot measuring the number of resulting clusters and the fraction of original genes retained (orange line corresponding orange Y axis). B) Boxplots showing average edge segregation of all clusters, indicating overall modularity of the network within each *T_d_*. C) A plot showing number of clusters enriched with at least one BP term and the total number of BP terms retained (orange line corresponding orange Y axis) and D) Boxplots summarizing the enrichment scores [−1*log(FDR)] of the hypergeometric p-values obtained by BP-cluster overlap analysis.

Overall, the network lost its stability and collapsed at *T_d_* values exceeding 0.80, as indicated by all measured parameters (Fig. 2). Hence, 1563 dense clusters obtained at *T_d_* 0.80 were tagged with functional (GO BP terms, KEGG pathways; Supplemental Data S2) and regulatory attributes (known *cis*-regulatory elements) by enrichment analysis (see Methods; Supplemental Data S3). We also mapped the expression patterns of each module spatially and temporally (seed compartment wise and maturation stage wise), by averaging the expression of module genes in each seed-compartment irrespective of the developmental stage or within each developmental stage irrespective of the seed compartment.

Visualizing some of these clusters in Cytoscape (Kohl et al., 2011) along with the functional attributes revealed a few remarkable properties of the seed network. The correspondence between the expression pattern of the cluster and the biological process (function) it represents match almost accurately in terms of the current knowledge in the literature. For example, one would expect the term ‘flavonoid metabolism’ enriched in clusters tagged with ‘seed coat’, the term ‘lipid storage’ and ‘photosynthesis’ enriched in clusters tagged with ‘endosperm’, and the term ‘auxin transport’ in clusters tagged with ‘embryos’. Not only were these terms correctly recovered by enrichment analysis, the observed inter- and intra-module connectivity between clusters was also as expected (Fig. 3). These observations reveal that the seed network is indeed modular, as sets of functionally related gene clusters or ‘modules’ get activated during different stages of seed development and in different seed compartments, and this nature of the seed network is accurately captured with the used clustering parameters.

**Figure 3:**
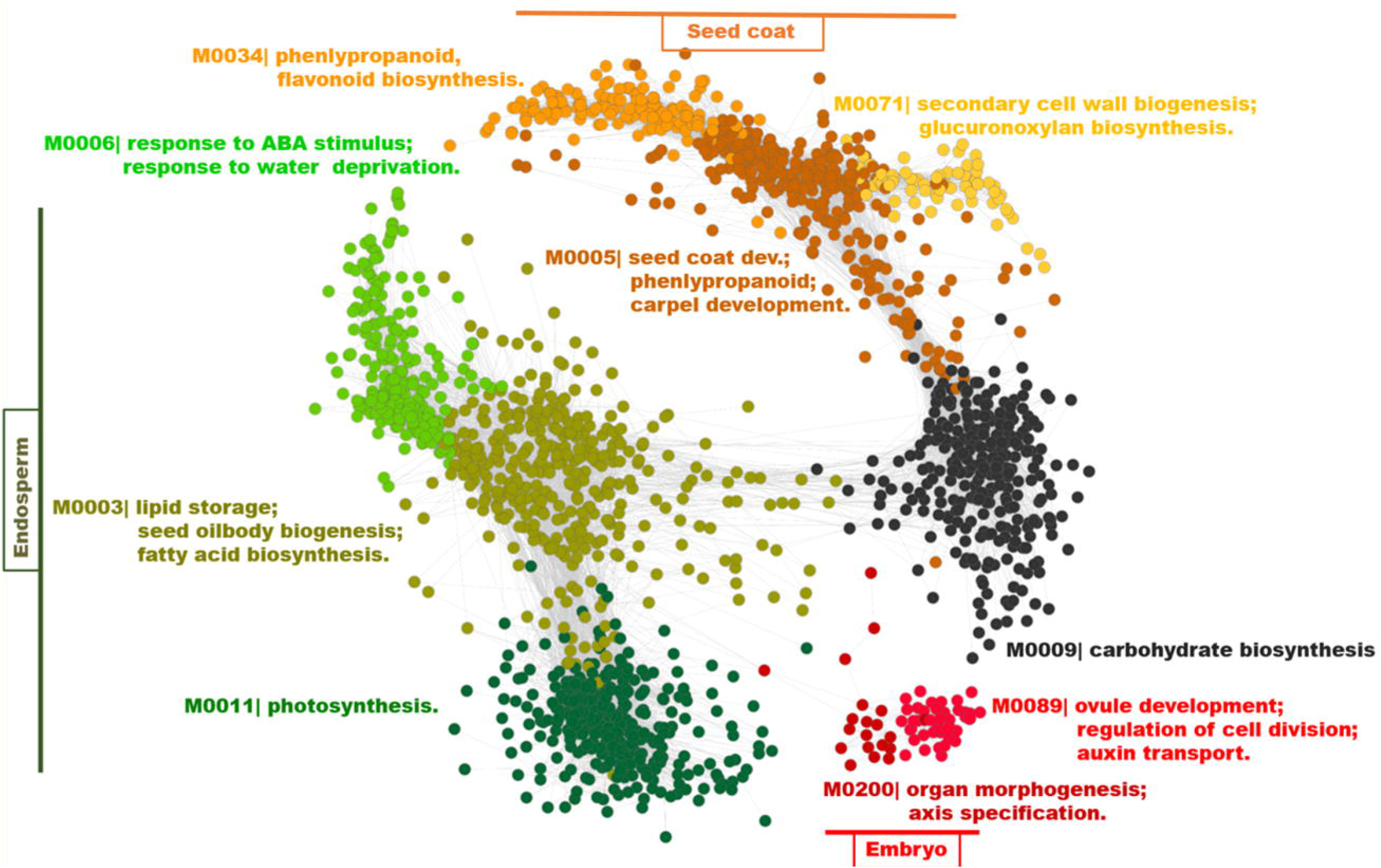
Visualization of seed modules. A graphical representation of seed coexpression modules was rendered using Cytoscape V3.3. Each circle represents a gene. Each module is uniquely color coded. Modules are grouped according to the seed compartments (indicated by horizontal or vertical lines and text boxes) and labelled with the BP term most highly overrepresented within each module. Genes are left unlabeled to facilitate visualization.

### Identifying transcriptional regulators of seed modules

Coexpression networks are usually devoid of information about gene regulation mediated by TFs. Simply classifying genes as targets of the TFs based on raw correlation scores is prone to occurrence of false positives. TFs can have high coexpression scores (Pearson’s correlation in this study) with a large fraction of genes in the genome due to indirect regulation. It thus becomes important to either identify and eliminate possible indirect targets or estimate how much an observed TF-target coexpression score ‘stands out’ within the background. The next step in our analysis was to incorporate a layer of regulatory information onto the seed coexpression network in a manner that utilizes an extensive data-driven weighted-analysis framework.

We scanned several Arabidopsis databases to identify a total of 1921 genes encoding TFs in Arabidopsis. We then transformed the normalized Pearson’s Correlation (PC) score between each TF-target gene pair into a ‘specific’ correlation score using the Context Likelihood of Relatedness (CLR) algorithm (Faith et al., 2007). This technique essentially down-weighs a TF-target gene pair with high PC score if the background distribution of PC scores of each gene in the pair is also high (see Methods; Supplemental Data S4). We then used these genome-wide specific correlation scores of each TF in a parametric analysis framework for gene set enrichment using the seed coexpression modules we defined in the preceding section as gene sets (Kim and Volsky, 2005). This step identified seed modules that contain (are enriched for) the most likely (high weighted) targets of TFs, essentially linking TFs to functional processes represented by the modules during different stages of seed development and in different seed compartments. It is important to note that no arbitrary cutoff score was chosen for selecting targets of each TF prior to enrichment analysis with modules. Instead, the entire genome was used, where each gene was a likely target weighted by the specific correlation score suggested by the underlying context of the expression dataset used. This unbiased approach allowed to bypass the estimation of cost-benefit ratio in selection of an optimal threshold.

The resulting seed-specific functional regulatory network is referred to as the ‘Seed Active Network’ or SANe. The regulatory core of SANe is essentially a bipartite graph represented as a matrix of 1819 TF regulators in rows and modules in columns, with each cell in the matrix representing a TF-module association score, and each module in turn tagged with functional (GO BP, KEGG and CYC pathways) and regulatory (TF and *cis*-regulatory elements) attributes, as well as spatial (seed compartment) and temporal (development phase) expression patterns (Fig. 4A). Visualizing the regulatory core of SANe reveals a dense interconnected network of TFs that associate with multiple modules suggesting a high redundancy or coordinate regulation of seed biological processes (Fig. 4B). The top 5 predicted regulators of a few selected modules (described later) revealed several known seed TFs reported in the literature while many others have been reported to play a role during other reproductive stages of the plant (Fig. 5; Supplemental Data S5).

**Figure 4:**
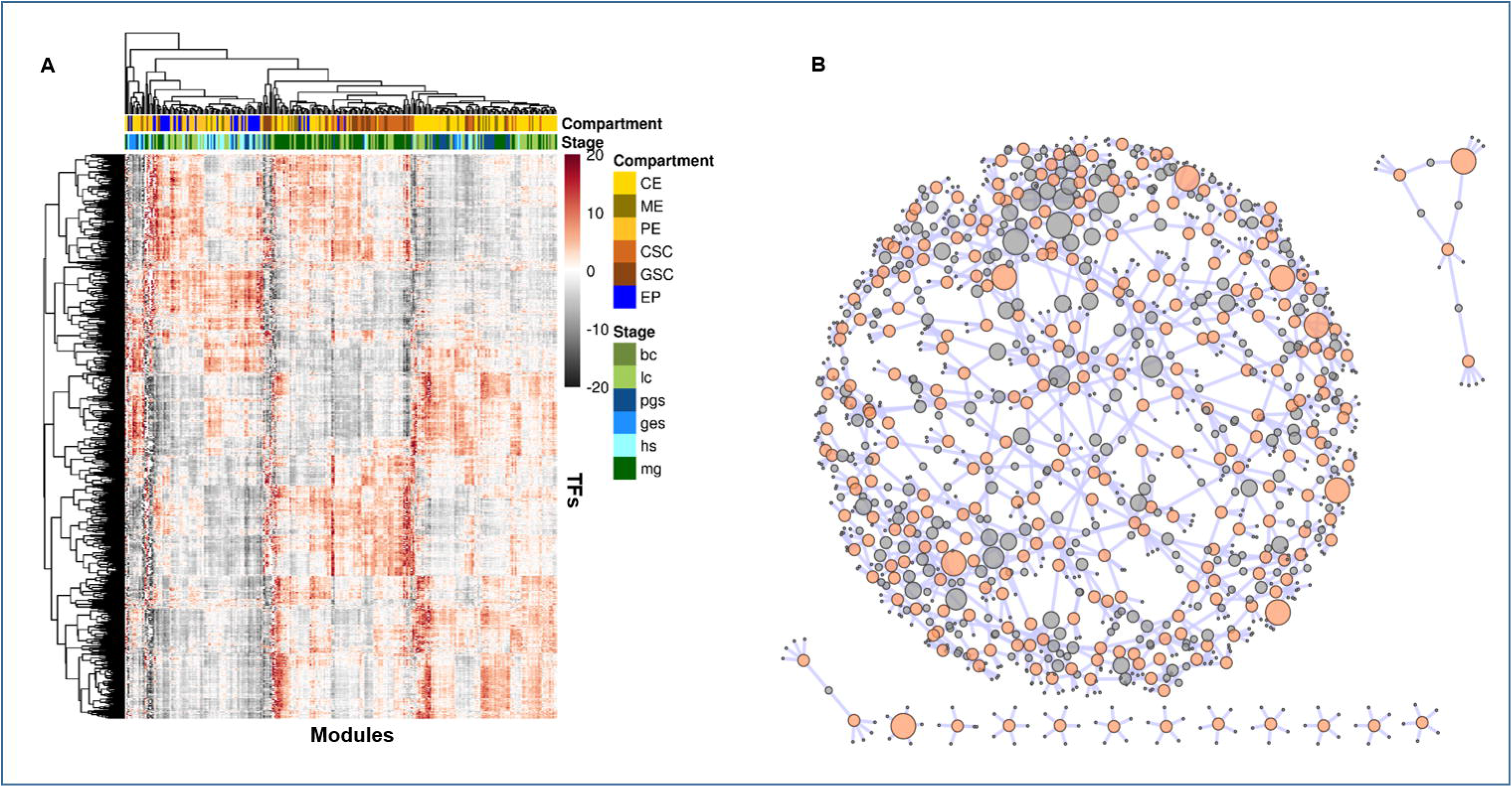
TF-Module Association Network. A) Heatmap representing association scores of 1819 TFs as regulators in the rows, and 10526 genes grouped into 278 coexpression modules represented along the columns. Each grid in the heatmap is color coded according to the level of enrichment of predicted targets of each TF regulator in the corresponding module. The red gradient indicates a positive score and grey indicates a negative score, estimated using the PAGE algorithm. Each seed compartment, in which the module with genes showing maximum expression, is color-coded and represented on top of the heatmap (first row), where CE is chalazal endosperm, ME is micropylar endosperm, PE is peripheral endosperm, CSC and GSC is chalazal and general seed coat, respectively, and EP is embryo proper. Each developmental stage, in which the module with genes showing maximum expression, maximum expression, is color-coded and represented on top of the heatmap (second row), where bc is bending cotyledon, lc is linear cotyledon, pgs is pre-globular, ges is globular, hs is heart and mg is mature green embryo stages. B) Predictions for each of the 278 modules were ranked and the top 5 predicted regulators for each module were visualized as a network graph. Each grey circle in the network plot is a TF and each orange circle is a module. The size of the grey circle is proportional to the out-going degree of the TF. Size of the orange circle was set to a constant, except for 9 bigger circles showing the modules described later in the main text. The network was visualized using Cytoscape version 3.3.0. Node names are hidden for ease in visualization. The Cytoscape sessions file is provided as supplemental data S3, which can be loaded into Cytoscape for node names and further exploration of the network. The heatmap was drawn using gplots package in the R statistical computing environment.

**Figure 5:**
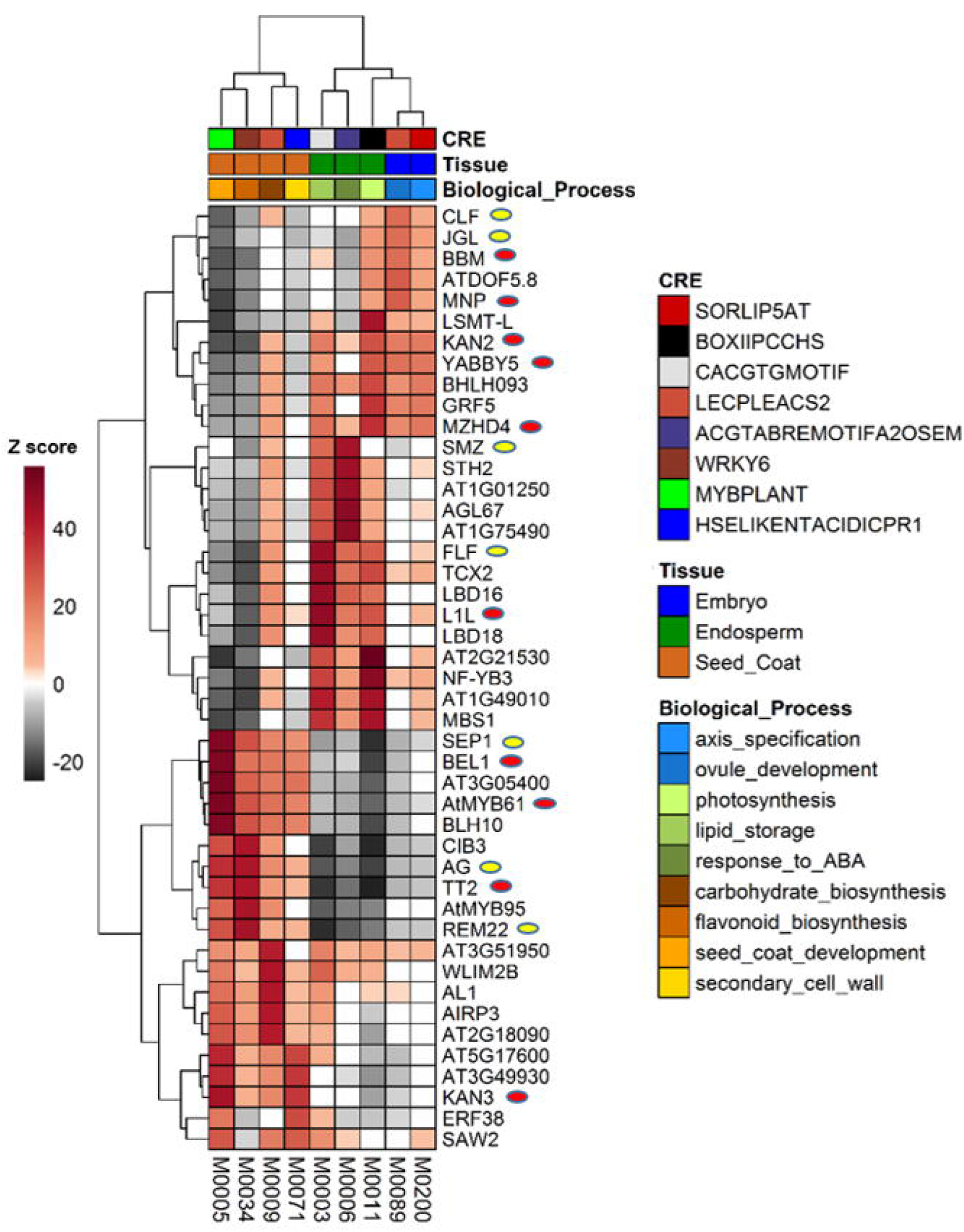
Heatmap representation of a subset of the SANe. Modules with high relative expression in embryo, endosperm and seed coat regions were extracted from SANe. Modules are shown in columns and for each module, the top 5 predicted TF regulators are shown in rows. Each grid in the heatmap is colored according to the association score estimated using the PAGE algorithm. Positive and negative scores are shaded in red or black gradient, respectively, as indicated by the color key. Literature identified TFs with validated seed-specific phenotypes or phenotypes observed in other reproductive stages/tissues are marked with a red ellipse or a yellow ellipse, respectively. CRE, cell-type and functional annotations for each module are shown above the heatmap (top three rows; colored boxes). Modules annotated for embryo, endosperm and seed coat are indicated in blue, green and brown boxes, respectively, in the middle row. CRE and functional annotation for each module are color-coded in the top and bottom rows, respectively.

### Estimation of differential regulation prioritizes hubs of the seed network

One important reason to model regulatory networks in a context-specific manner is that the resulting data lends itself for comparison of different biological contexts to understand the dynamics of transcriptional regulatory networks. Here, we attempted to estimate differences in the targeting patterns of TFs, or network rewiring in this tissue-specific network compared to a ‘global’ network not specific to any specific tissue/organ of the plant.

To make such a comparison, we created another contextually distinct expression compendium with gene expression values from various other organs of the Arabidopsis plant, including vegetative and seedling growth stages. We followed the same steps of associating TF to seed modules (only the steps in top-left box with orange dashed lines in Fig. 1), but this time using TF-target specific correlation scores from this global expression dataset to create the global network. We then quantified the differences in module association scores for each TF between this global network and the seed-specific network. TFs were then ranked on the basis of the magnitude of this differential score of association with all seed modules (see Methods; Supplemental Data S6). Remarkably, LEC1, a well-characterized master regulator of seed development (Kagaya et al., 2005) was found at rank #1, indicating that LEC1 has the most rewired connections during seed development. Furthermore, other members of the LAFL network (Jia et al., 2013) were recovered at the top of all rankings, with ABI3 at rank #50, FUS3 at rank #8 and LEC2 at rank #7 (the range of ranks was from 1 to 1819; see Methods). We observed that the top associated modules with these TFs represent similar sets of physiological processes and in many cases the direction of regulation was found opposite between the two networks compared (Fig. 6). For example, while LEC2 maintained a strong positive association with photosynthesis, ribosome biogenesis and RNA metabolism in the seed network (Fig. 6A), its association with these processes in the global network appear to be broken as indicated with a negative association score (Fig. 6B). The association of ABI3 with the GO BP term ‘response to abscisic acid stimulus’ was observed to be stronger in the global network than the seed network (Fig. 6C and 6D). FUS3 was found positively associated with photosynthesis and steroid biosynthesis only in the seed network (Fig. 6E and 6F), while L1L was found to positively regulate DNA methylation only in the seed network (Fig. 6G and 6H).

**Figure 6:**
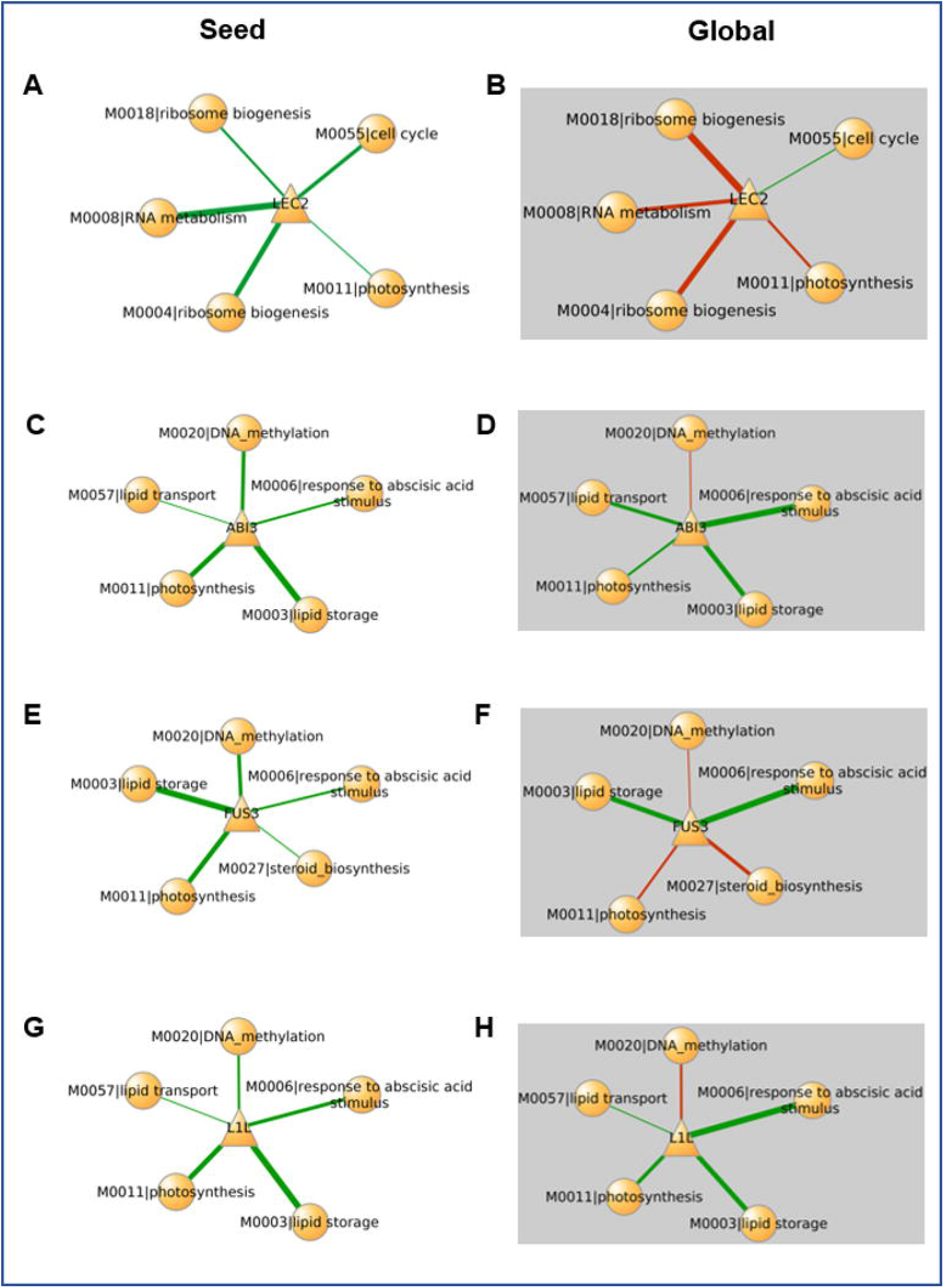
Differential rewiring of TFs in the seed network. The association TFs with seed modules was estimated in networks built around two different biological contexts: the Seed network (left panel) and a global network (right panel with grey boxes). TFs (center triangle-shaped nodes) were connected to seed coexpression modules (circular peripheral nodes) if the predicted targets of corresponding TF are enriched in the module, tested using parametric analysis of geneset enrichment. Green and red edges indicate positive and negative enrichment score, respectively. A) and B) show association of LEC2 in seed network and global network, respectively. C) and D) show association of ABI3, E) and F) show association of FUS3 and G) and H) show association of LEC1-like.

To evaluate these TF rankings more comprehensively and on a genome-wide scale, we identified Arabidopsis TFs that have evidence of seed-related phenotypes upon genetic intervention (over expression or knockout) in the literature. Out of 63 such TFs reported, 43 were part of our list of TFs used for building the network (Supplementary Data S7). We used the decile enrichment test (Krishnan et al., 2016) to evaluate whether a larger fraction of TFs from this literature-curated Validated Gene Set (VGS) were present in our first decile (top 10% of all rankings; see Methods). The test revealed a significant enrichment of VGS in our first decile rankings (binomial test *p*=3.67×10^−6^; Fig. 7A). The enrichment also holds significance for experimentally confirmed targets of TFs in VGS (*p*=1.544×10^−5^; Fig. 7B). Overall, these results indicated that our strategy of prioritizing seed-specific TFs is accurate and makes other understudied TFs within the first decile as the primary candidates for reconstruction the seed development network experimentally.

**Figure 7:**
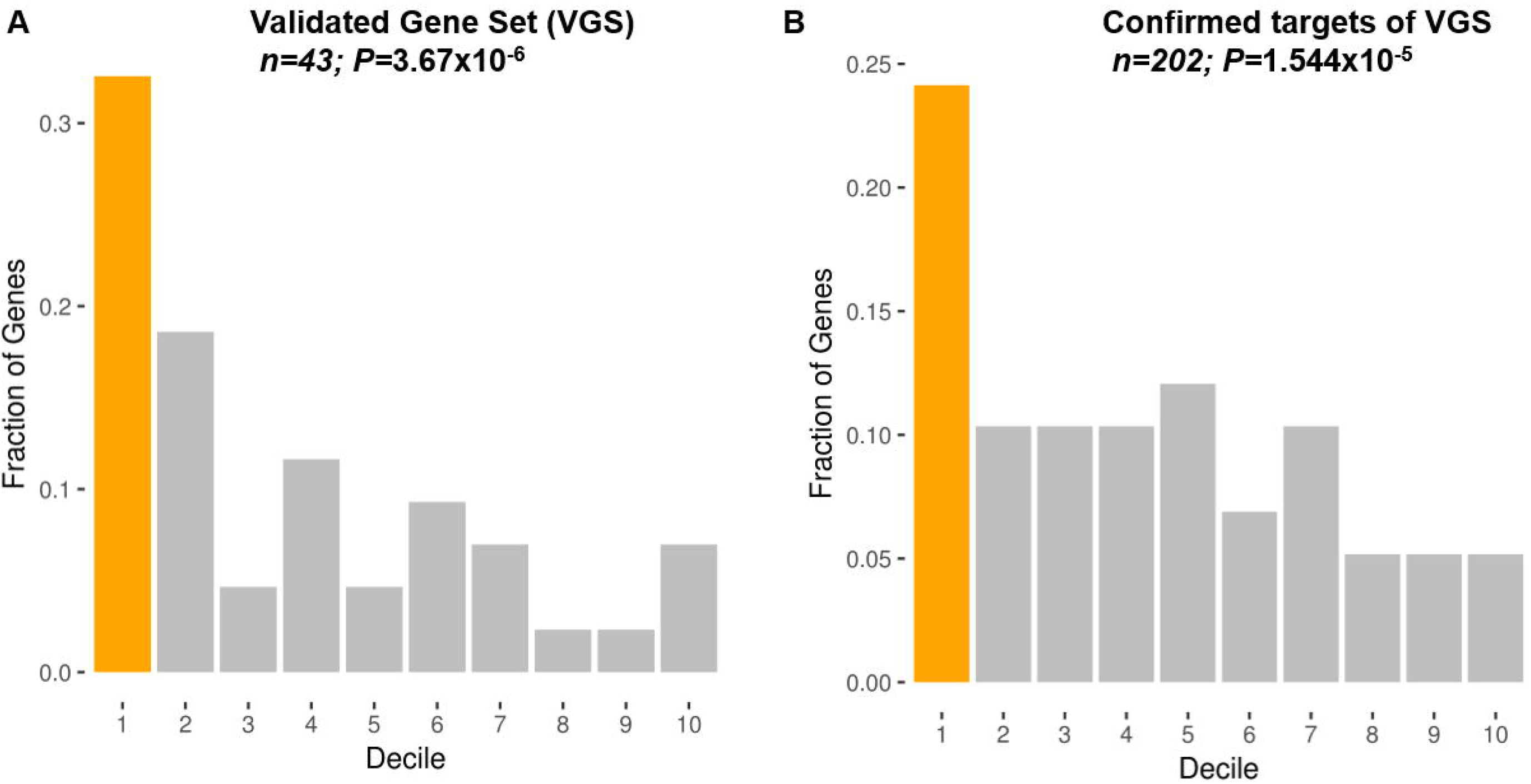
Decile Enrichment Test. TFs that significantly changed association with seed modules between the seed-specific network and the global network were identified and ranked according to the differential regulation score. Total rankings (1 to 1819) was divided into 10 equal parts (deciles) and the first decile tested for enrichment of A) literature curated list of TFs that have experimentally confirmed morphological, physiological, and/or metabolic phenotypes during seed development and B) experimentally validated targets of these TFs identified from the AGRIS server. The first decile in both the plots are shown in orange and the number of genes in each tested gene set is shown on the top of each plot along with the *p* values from the binomial test.

### Using the SANe webtool to survey seed genes

We created a web interface to provide easy access to the data generated in building SANe. The platform is hosted at https://plantstress-pereira.uark.edu/SANe/ and presents users with tools to select specific modules of interest, predict functional regulons of a TF or perform gene set enrichment analysis using seed modules. The module enrichment tool serves as an alternative to the traditional pathway/GO BP enrichment analysis of a differential expression test and provides additional information about the regulators likely involved in the observed phenotype in the experiment. For example, uploading the differentially expressed transcriptome of FUS3, LEC1 and LEC2 mutants (GSE61686) into the cluster enrichment tool of SANe revealed repression of modules M0003 and M0011 representing lipid storage and photosynthesis, respectively, in all the three mutants (Supplemental Data S8). SANe can be used to investigate different stages and compartments of seed development and generate testable hypotheses. We describe a few key modules identified from SANe and demonstrate the usage of the webserver below.

### Modules for early embryo development

Three modules designated as M0089, M0200 and M0277 comprised 54, 31 and 33 genes, respectively, expressed at relatively high levels in the embryonic tissue when compared to other seed compartments (Fig. 8A). These genes are significantly enriched with BP terms such as “organ development”, “tissue development”, “axis specification” and “auxin transport”. This is consistent with processes related to embryo development, involving morphogenesis-related and other cellular processes that govern gene activity related to cell division and expansion, maintenance of meristems and cell fate determination (Wendrich and Weijers, 2013).

**Figure 8:**
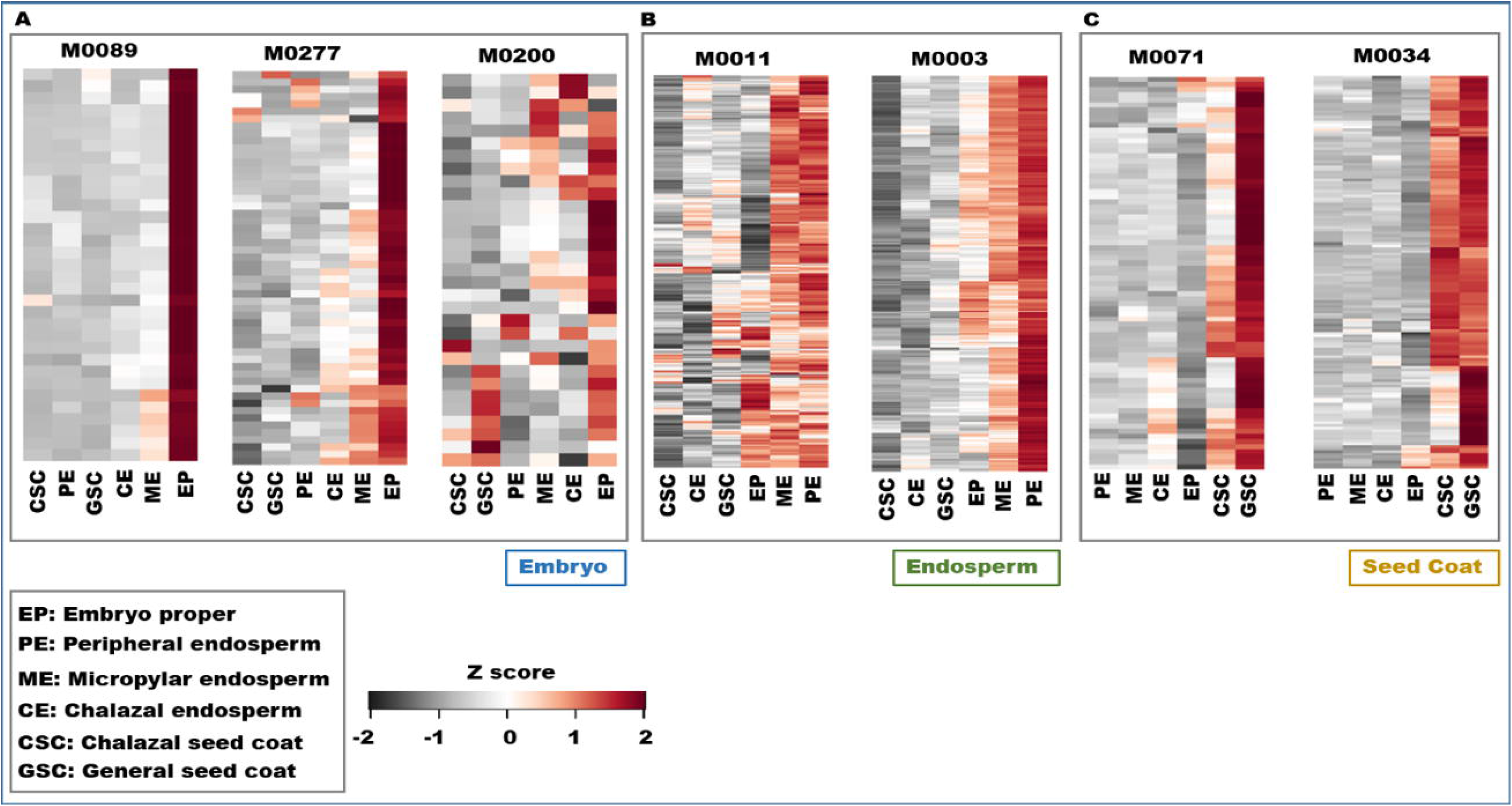
Expression profiling of gene modules. Expression patterns of modules in embryo, endosperm and seed coat regions represented as heatmaps in A), B), and C), respectively. Seed compartments are represented as columns and genes as rows. Gene names are hidden for ease in visualization. Expression values of genes in each module were averaged across samples from the same tissue-type/seed-compartment (embryo, endosperm and seed coat). Average expression values were scaled and represented as Z scores in the heatmaps. Red indicates high expression of a gene in a particular compartment and black gradient indicates low expression relative to other compartments.

**Figure 9:**
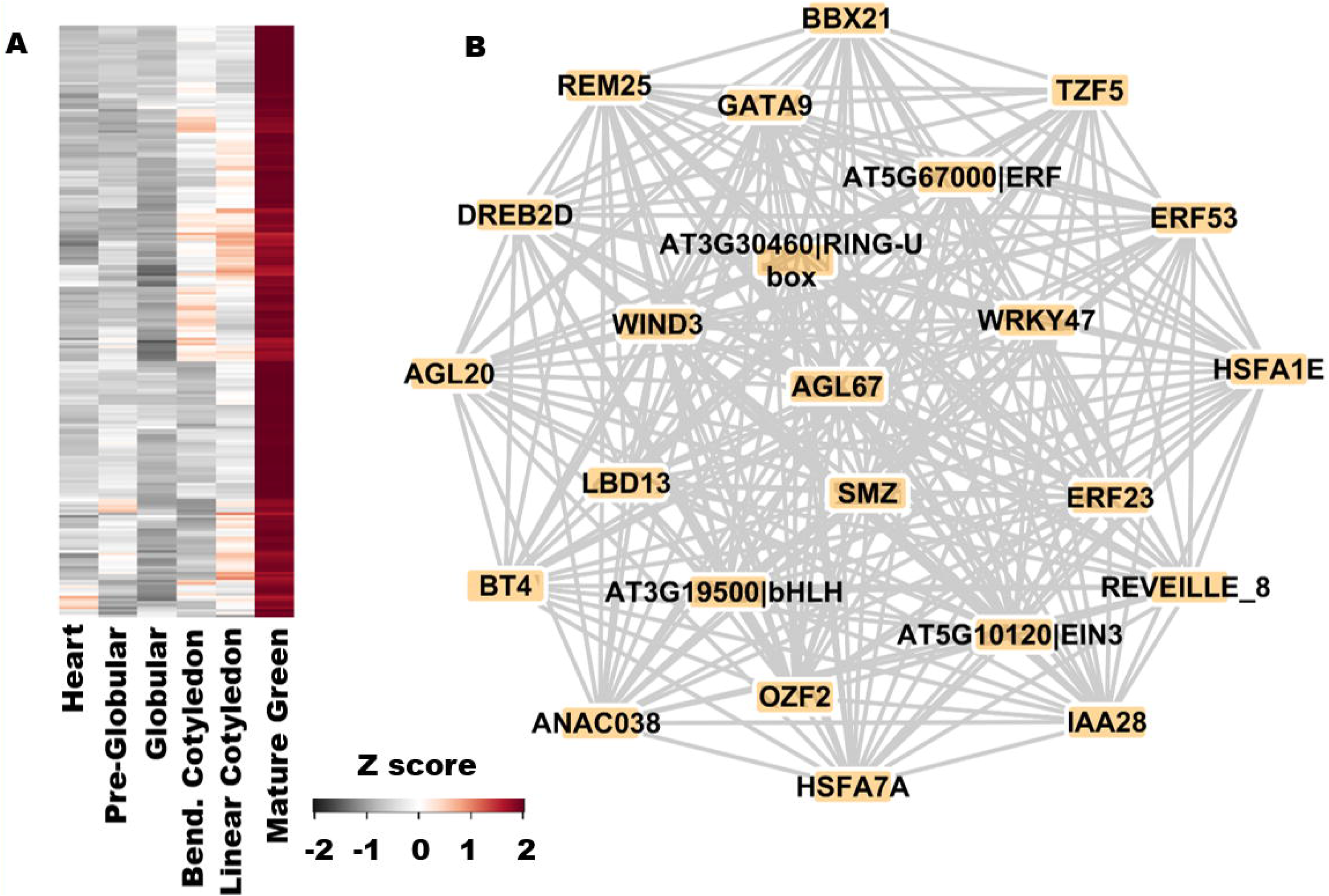
Module M0006. A) Expression patterns of genes in module M0006. Seed developmental stages are represented as columns and genes as rows. Gene names are hidden for ease in visualization. B) Coexpression links between TFs in M0006. Nodes are labeled according to their corresponding gene symbols if present in TAIR, or labeled with their corresponding locus ID and the family they belong.

M0089 harbors genes related to reproductive tissue development and cell division. ATDOF5.8 (AT5G66940) was predicted as the top regulator of M0089. The *ATDOF5.8* gene is most highly expressed in embryo and meristem cells (Supplemental Fig. 1A) based on the Genevisible tool in GENEVESTIGATOR (Zimmermann et al., 2004). It has been shown that ATFOD5.8 is an abiotic stress-related TF that acts upstream of ANAC069/NTM2 (AT4G01550) (He et al., 2015). Interestingly, the *NTM2* gene resides at a locus adjacent to another NAC domain TF, NTM1 (AT4G01540), a regulator of cell division in vegetative tissues (Kim et al., 2006). Kim et al. did not detect *NTM2* expression in leaves by RT-PCR. However, they indicated that because both *NTM* genes have similar structural organization, encoding proteins with a few differences in the protein chain, NTM2 could be involved in similar processes in other tissues. Our predictions suggest that *NTM2* could be in the ATDOF5.8 regulon associated with modulating cell division activity in the seed. This leads to a new testable hypothesis pertaining to regulation of cell division during embryogenesis. Among other known regulators, BABY BOOM (BBM, AT5G17430) was predicted as one of the top ranked TFs (rank 4) of M0089. BBM is an AP2 TF that regulates the embryonic phase of development (Boutilier et al., 2002).

YAB5 (AT2G26580) and ATMYB62 (AT1G68320) were predicted the top ranked regulators of M0200 and M0277, respectively. While the agreement of YAB5 as a determinant of abaxial leaf polarity (Husbands et al., 2015) and enrichment of M0200 with GO BP term “axis specification” (GO:0009798) justifies this association, the association of ATMYB62 with M0277 indicates a hormonal interaction likely representing a transition between the growth stages. *ATMYB62* encodes a regulator of gibberellic acid biosynthesis (Devaiah et al., 2009) and is expressed specifically during seed development (Belmonte et al., 2013). M0277 is enriched with “auxin transport” genes (GO:0009926). The *ATMYB62* gene is preferentially expressed in the abscission zone and other reproductive tissues (Supplemental Fig. 1B).

### Modules for Endosperm Development

The endosperm has a profound influence on seed development by supplying nutrients to the growing embryo (Portereiko et al., 2006; Chen et al., 2015). The importance of endosperm cellularization for embryo vitality has been shown through mutants deficient in endosperm-specific fertilization events (Kohler et al., 2003). The overall seed size depends on endosperm development and is controlled through the relative dosage of accumulated paternal and maternal alleles (Luo et al., 2005).

We found that genes in modules M0003 and M0011 had maximal expression levels in endosperm tissues (Fig. 8B). M0003 is significantly enriched with genes involved in lipid storage (GO:0019915) and fatty acid biosynthesis (GO:0006633). LEC1-LIKE (L1L, AT5G47670) emerged as the top regulator of this module. LIL is related to LEAFY COTYLEDON 1 (LEC1) and functions during early seed filling as a positive regulator of seed storage compound accumulation (Kwong et al., 2003). Interestingly, L1L is also part of this module indicating that, apart from being a master regulator, its activity is also modulated during the late seed filling stages as observed previously (Kwong et al., 2003), which correlates with the overall expression pattern of genes within this module (Supplemental Fig. S2). The presence of 44 other TFs in this module, including FUS3 and ABI3, key regulators of seed maturation (Keith et al., 1994; Luerßen et al., 1998; Yamamoto et al., 2009), points to the importance of this module in nutrient supply to the developing embryo. LDB18 (AT2G45420) is a LOB-domain containing protein of unknown function predicted as the second ranked regulator of this module. GENEVESTIGATOR analysis showed that both *L1L* and *LDB18* are most highly expressed in the micropylar endosperm (Supplemental Fig. S3).

M0011 is comprised of 357 genes including 7 TFs and is characterized by containing genes with high expression levels in the micropylar endosperm (ME) and the peripheral endosperm (PE). GO enrichment analysis showed the highest scores for photosynthetic genes (GO:0015979) in this module. Close examination of these genes revealed that virtually all aspects associated with chloroplast formation and function were represented, including chloroplast biogenesis and membrane component synthesis, chlorophyll biosynthesis, plastidic gene expression, photosynthetic light harvesting and electron transport chain, ATP production, redox regulation and oxidative stress responses, Calvin cycle and photosynthetic metabolism, metabolite transport, and retrograde signaling. Interestingly, genes encoding photorespiratory enzymes (glycine decarboxylase, glyoxylate reductase, and hydroxypyruvate reductase) were also present in M0011. Developing oilseeds are known to keep extremely high levels of CO2 that would suppress photorespiration (Goffman et al., 2004), and the implications of expression of these genes on photosynthetic metabolism are not clear.

The presence of mostly photosynthetic genes in M0011 seems also unusual, but the results are consistent with findings of (Belmonte et al., 2013), showing that specific types of endosperm cells are photosynthetic, as they contain differentiated chloroplasts and express photosynthesis-related genes. Fully differentiated embryos at the seed-filling stages and the chlorophyll-containing inner integument ii2 of the seed coat are parts of oilseeds that are also capable of photosynthesis (Belmonte et al., 2013; Sreenivasulu and Wobus, 2013). Although seeds obtain the majority of nutrients maternally, Arabidopsis embryos remain green during seed filling and maintain a functional photosynthesis apparatus similar to that in leaves (Allorent et al., 2015). As part of photoheterotrophic metabolism, photosynthesis provides at least 50% of reductant in oilseed embryos and CO2 is re-fixed through the Rubisco bypass that helps to increase carbon-use efficiency in developing oilseeds (Ruuska et al., 2004; Schwender et al., 2004; Goffman et al., 2005; Fait et al., 2006). The roles for photosynthesis in ME and PE remain to be investigated and include (i) providing carbon and energy for storage compound accumulation in the endosperm and the embryo and (ii) increasing the availability of oxygen to the endosperm and differentiating, yet-to-be photosynthetic, embryos in a high-CO2 environment.

CRE analysis revealed the highest number of motifs enriched in the promoters of genes in M0011, suggesting extensive coordination between different regulators. Light-related motifs BOXIIPCCHS (ACGTGGC), IRO2OS (CACGTGG3), IBOXCORENT (GATAAGR) and the ABA-responsive element ACGTABREMOTIFA2OSEM are the most over-represented motifs in this module. The highest ranked regulator of M0011 is a SMAD/FHA domain-containing protein (AT2G21530) that is most highly expressed in the cotyledons (Supplemental Fig. S4A). The known seed-specific regulator of oil synthesis and accumulation WRI1 (AT3G54320) was identified as the sixth ranked regulator of this module and is suggested to be predominantly expressed in the embryo and endosperm (Supplemental Fig. S4B). *WRI1* encodes an AP2/ERF-binding protein and *wri1* seeds have about 80% reduction in oil content relative to the wild type seeds (Ruuska et al., 2002). Genetic and molecular analysis revealed that WRI1 functions downstream of LEC1 (Baud et al., 2007). Along with WRI1 itself, six other TFs are part of this module, including AT2G21530, a zinc finger (C2H2) protein (AT3G02970), NF-YB3 (AT4G14540), PLT3 (AT5G10510), GIF1 (AT5G28640) and PLT7 (AT5G65510).

### Modules for Seed Coat development

The seed coat has important functions in protecting the embryo from pathogen attack and mechanical stress. The seed coat encases the dormant seed until germination and maintains the dehydrated state by being impermeable to water. M0034 is comprised of 149 genes with the highest expression in general, and specifically in chalazal seed coat relative to other tissues (Fig. 8C). This module is enriched with genes annotated under the GO BP terms “phenylpropanoid biosynthetic process” (GO:0009699) and “flavonoid biosynthesis process” (GO: 0009813). The AP2/B3-like TF AT3G46770 is highly expressed in seed coat (Supplemental Fig. S5A) and predicted as the top regulator in this module. B3 domain TFs are well known for functioning during seed development and transition into dormancy in *Arabidopsis* (Suzuki and McCarty, 2008) and, to some extent, their functions are conserved in cereals (Grimault et al., 2015). The seed-coat-specific expression of AT3G46770 is a compelling incentive for testing AT3G46770 mutants for seed-related phenotypes, which to the best of our knowledge, has never been considered. There were 21 other TFs belonging to this module, of which six are part of the MYB family. TRANSPARENT TESTA 2 (TT2), a MYB family regulator of flavonoid synthesis (Nesi et al., 2001), was ranked fourth in our predictions for this module.

M0071 is composed of 77 genes encoding, surprisingly, only 3 TFs, ERF38 (AT2G35700), BEL1-LIKE HOMEODOMAIN 1 (BLH1, AT2G35940) and a C2H2 super family protein (AT3G49930). This module is enriched with genes involved in “xylan metabolic process” (GO:0045491), “cell wall biogenesis” (GO:0009834), and “carbohydrate biosynthetic process” (GO:0016051). KANADI3/KAN3 (AT4G17695) was predicted as the top regulator of this module. KANADI group of functionally redundant TFs (KAN1, 2, and 3) has been shown to play roles in modulating auxin signaling during embryogenesis and organ polarity (Eshed et al., 2004; McAbee et al., 2006; Izhaki and Bowman, 2007). In the case of another KANADI TF, KAN4, encoded by the *ABERRANT TESTA SHAPE* gene, the lack of the KAN4 protein resulted in congenital integument fusion (McAbee et al., 2006). It is reasonable to hypothesize that KAN3 could be acting in a redundant manner with KAN4 to regulate seed coat formation during late stages of maturation, as the expression pattern of *KAN3* is higher in seed coat than in other organs or cell types (Supplemental Fig. S5B).

### The desiccation tolerance module (M0006) is regulated by AGL67

M0006 is comprised of 220 genes expressed predominantly during the mature green stage (Fig. 9A), and enriched with genes involved in “response to abscisic acid stimulus” (GO:0009737), “response to water” (GO:0009415) and terms related to embryonic development (GO:0009793), altogether suggesting an involvement of these genes in acquisition of desiccation tolerance (DT). We predicted AGL67 (AT1G77950) as a major regulator of this module, among 23 other TFs that are part of this module (Fig. 9B). AGL67 has been recently confirmed as a major TF involved in acquisition of DT (González-Morales et al., 2016), validating our prediction. Additionally, the authors of this study analyzed the mutants of 16 genes (TFs and non-TFs) that had reduced germination percentage, of which 12 are in our network and 7 of these are a part of M0006. These 7 genes include PIRL8 (AT4G26050), ERF23 (AT1G01250), OBAP1A (AT1G05510), DREB2D (AT1G75490), AT1G77950 (AGL67), AT2G19320 and MSRB6 (AT4G04840).

### Experimental validation of SANe predictions

A few first decile TF predictions from SANe were tested by reverse genetics and analytical methods to determine if these TFs are involved in different aspects of seed development and metabolism. In some cases, homozygous mutants could not be obtained likely due to potential or known embryo lethality (Supplemental Data S9). Mutants in TFs known to be involved in seed development and metabolism, such as in bZIP67 (AT3G44460), L1L (AT5G47670), and MYB118 (AT3G27785) were included as controls and their dry seeds showed altered fatty acid composition. For example, MYB118 is known to repress endosperm maturation and oil biosynthesis through suppression of WRINKLED1 (WRI1) and oil biosynthetic genes in the endosperm (Barthole et al., 2014; Fatihi et al., 2016). However, together with MYB115, it activates fatty acid desaturases involved in the formation of omega-7 fatty acids (Troncoso-Ponce et al., 2016). Dry seeds of the *myb118* mutant (Salk_111812) showed decreased levels of C18:1Δ7 (*p* < 0.01) and elevated levels of C20:0 and C22:1 (*p* < 0.02), but oil content remained unchanged (Supplemental Data S10).

TZF6/PEI1 (AT5G07500) is involved in the formation of heart-stage embryos (Li and Thomas, 1998). SANe suggested TZF6 to be significantly associated with M0003 involved in fatty acid metabolism. The corresponding Salk_101961 mutant showed significantly (*p* < 0.03) reduced levels of several fatty acids, including C18:1Δ7 (66% of the wild type levels), C18:3 (75%), C20:1 (75%), and C22:1 (81%) in dry seeds, though the total oil content remained similar to that of the wild type. A 30% decrease in seed storage protein content (*p* < 0.005) was observed in Salk_079930. This mutant is disrupted in the AT5G07160 gene encoding a seed-specific bZIP TF of previously unknown function and predicted by SANe to be a regulator of M0002 enriched with genes that function in the phenylpropanoid pathway. The specific function of AT5G07160 is not known, but there is a connection between FUS3 (positive regulator of protein synthesis in the seed) and the global regulators of phenylpropanoid synthesis TT2 and TRANSPARENT TESTA GLABRA1 (TTG1). FUS3 is a direct negative regulator of TTG1 and seed carbon partitioning and seed storage protein and oil synthesis are dependent on the phosphorylation state of TTG1, which determines its ability to physically interact with TT2 (Chen et al., 2015; Li et al., 2018). Collectively, seed storage proteins and oil encompass about 70 – 80% of seed dry biomass. C18:3 and C20:1 are abundant in Arabidopsis seed oil. As such, 20 – 30% alterations to seed storage compound composition represents significant alterations in oil and protein levels.

Principal Component Analysis (PCA) on correlations of metabolite levels was used to assess if any of the analyzed mutants had significantly different metabolomes in dry seeds. None of the mutants were different from the wild type from the global perspective (Supplemental Figure 6), though many small, but statistically significant changes (< 3 fold, *p* < 0.02) in metabolite levels were observed between wild type and mutant seeds (Supplemental Data S10). For example, seed Ser levels were doubled in the *lec1* mutant compared to the wild type. In contrast, Asn levels were reduced to 36, 40, and 42% of the wild type levels in the *MINISEED 3* (*MINI3*, At1g55600), *NUCLEAR FACTOR-YC7* (*NF-YC7*, At5g50470), and Salk_014315 (AT3G10590) mutants, respectively. While MINI3 positively regulates seed size (Luo et al., 2005; Meng et al., 2016), functions of the other two TFs were previously unknown. SANe correctly predicted *NF-YC7* as the regulator of M0030 involved in amino-acid transport, and AT3G10590 as the regulator of M0022 involved in trehalose biosynthesis.

In contrast to seed storage compounds, metabolite levels change rapidly (within seconds and minutes) depending on environmental conditions and only large statistically significant changes in metabolite levels should be considered. A 3.3-fold increase in galactinol levels were observed in the Salk_085497 mutant in the AT3G44460 gene encoding bZIP67 that was predicted (and is known) to be a TF involved in fatty acid metabolism (M0003). Galactinol accumulates during late maturation and seed desiccation and is associated with seed longevity(de Souza Vidigal et al., 2016; Li et al., 2017). This potential connection between fatty acid synthesis and galactinol remains to be explored. However, the most significant changes (≥ 3 fold, *p* < 0.01) were related to unknown sugars, and, as such, potential connections could not be predicted. The *hdg3* mutant (Salk_033462) in the homeobox-leucine zipper family TF HOMEODOMAIN GLABROUS 3 gene (AT2G32370) involved in cotyledon development (Nakamura et al., 2006) accumulated 6.6 and 31% of an unknown tri and disaccharide, respectively, relative to the wild type. Levels of another unknown trisaccharide were reduced to 7.4% of the wild type levels in the *lec1* mutant. The Salk_033801 mutant in the MADS-box TF of unknown function encoded by AT5G26630 accumulated 10% of the wild type levels of an unknown hexose. Disruption of the *MINI3* and *HEAT SHOCK TRANSCRIPTION FACTOR A9* (*HSFA9*, AT5G54070) genes resulted in 12 and 21%, respectively, of the wild type levels of an unknown disaccharide. HSFA9 is regulated by ABI3 and involved in late seed maturation (Kotak et al., 2007).

## Discussion

We previously observed that associating TFs to sets of biologically coherent genes (instead of individual genes) yields statistically strong functional predictions in the context of TFs in rice (Ambavaram et al., 2014). In order to accommodate predictions on a genome-wide scale and in a tissue-specific manner for the development of seed regulatory network, we made a few modifications to this experimentally validated network analysis method. This optimized new technique was used to build the Seed Active Network (SANe) that serves as a regulatory map of the process of seed development in the model Arabidopsis. SANe facilitated the identification of TFs that regulate functional modules of seed development, a layer of information typically not revealed by coexpression analysis.

The primary objective of this network analysis pipeline was to capture gene regulation information in a tissue-specific manner. To examine the validity of our approach and to identify the distinguishing characteristics of the seed regulatory network that differed from a global network (non-tissue specific regulatory network), we compared both networks and ranked TFs according to the magnitude of their ‘differential wiring’ in the seed. As expected, several known TFs necessary for seed development were found towards the top of the TF-wide ranking produced by our analysis. All known global regulators, LEC1, LEC1-like, LEC2, ABI and FUS3 were recovered within the top10% of all ranks. Since this method achieved a high recall, we reasoned that other top-predicted TFs – along with their coexpression neighborhood – will pave way to identification of transcriptional networks modulated specifically during seed development or involved in important seed functions. Along with other novel TFs that emerged as the top regulators of important modules, these TFs now become the primary candidates for testing seed phenotypes largely reducing the search space for experimental biologists.

It appears that during seed development, photosynthesis and storage compound synthesis is tightly coordinated by several regulators acting coordinately. This was evident from CRE enrichment analysis, as two complementary methods detected the module annotated for photosynthesis and related processes (M0011) harboring genes with the largest number of known plant motifs in their promoters when compared to the rest of the modules. Coordinate regulation of photosynthetic carbon metabolism has been shown previously (Bailey et al., 2007; Ambavaram et al., 2014). Our analysis reveals that much of the processes related to embryo development, such as cell division and differentiation, are conserved throughout the plant life cycle as observed by similar roles of regulatory genes in developing embryos and roots. However, plants have developed intrinsic mechanisms that can modulate gene activity in specialized cells, perhaps as duplicated genes with similar functional roles. Such a phenomenon was evident in the case of two TF genes, *NTM1* and *NTM2* that are in close proximity to each other and possibly have similar biological roles in distinct parts of a plant. The seed regulatory data generated by our work has the potential to further our knowledge of fundamental processes involved in diverse specific aspects of seed development, and can be extrapolated to related agriculturally important crops due to conservation of these basic processes (Magallón and Sanderson, 2002; Comparot-Moss and Denyer, 2009; Vriet et al., 2010).

The SANe webtool accompanying this paper can be used to identify modules with high expression in specific seed compartments, including embryo, endosperm and seed coat. We observed that, in most of the cases, the top predicted regulators of these modules are already known to be involved in seed development, self-validating our approach. Several additional regulators are known to modulate other processes, including flower development, indicating conserved regulons of pre-fertilization events. Our results suggest that associating regulators to gene sets with a shared function, as opposed to individual genes, provides biologically plausible predictions that are worth for validating *in planta* phenotypes using reverse genetics. The webtool is supported with query driven tools to enable a network-based discovery of seed regulatory mechanisms. Further, SANe will also help researchers align their newly generated seed transcriptomes to our network and identify TFs most relevant to their dataset. For example, we queried the transcriptomes of FUS3, LEC1 and LEC2 with the cluster enrichment tool of SANe, and found modules related to fatty-acids as the most highly perturbed, which corresponds to the known functional roles of these TFs (Supplemental Data S8).

Our network is different from other coexpression-based networks aimed at seed development on several counts. First, our method of analysis does not aim at predicting individual targets of TFs, instead SANe treats each gene in the genome as a potential target, weighted by the likelihood functional relatedness, thereby bypassing the requirement of target selection thresholds in regulatory network inference. Second, SANe significantly improves the interpretation of coexpression data by adding a layer of regulatory information on identified gene clusters, essentially exhausting the information content in the expression dataset. Third, SANe a genome-scale functional gene regulatory network covering a large fraction of the Arabidopsis genome, hence providing a starting point for scalable analyses.

The users of SANe should bear in mind that our regulatory network analysis method has been only tested for making functional predictions for TFs and might not accurately predict individual targets of a given TF. This limitation is partly due to the use of a single data-type; a heterogenous approach should be undertaken (e.g. high-throughput DNA binding essays in conjunction with expression data) for studies aiming at validation of specific individual targets. Nevertheless, the statistically significant functional associations predicted here are of superior quality, as seen in evidence from the literature, and can serve as the first step in selecting TFs for targeted downstream experiments. The network inference pipeline presented here can be used to enhance any expression network analysis-based study.

Based on our results, a cell- and developmental stage-specific network inference provides superior quality of predictions in the context of known information. Our network analysis pipeline can be further used to systematically increase this information-base for a variety of plant organs (e.g., parts from a post-germination stage network). Comparisons of different stage/tissue specific networks will throw light on the changing molecular mechanisms of a cell and reveal differentially modulated transcriptional networks during different growth stages.

### Experimental Procedures

#### Chemicals

All reagents and metabolite standards (analytical or higher purity) were obtained from Sigma-Aldrich (St. Louis, MO) or Thermo Fisher Scientific (Waltham, MA). Solvents used for metabolite extraction were MS grade. Multiply labeled internal standards were used as internal standards for gas chromatography-mass spectrometry (GC-MS) and were purchased from Cambridge Isotope Laboratories, Inc. (Tewksbury, MA). Internal standards included: (i) heptadecanoic acid for free and lipid-derived fatty acids, (ii) [U-^13^C_6_]-glucose for sugars, sugar alcohols, sugar acids, and sugar-phosphates, (iii) [2,2,4,4-D_4_]-citrate for carboxylic and other organic acids, and (iv) norvaline for amino acids and organic amines. These internal standards were added to samples prior to the extractions and used as described (Collakova et al., 2013; Schneider et al., 2016).

#### Gene expression quantification

Affymetrix ATH1 Arabidopsis gene expression data was downloaded from GEO, and 6 datasets were selected from the super series labeled GSE12404 for the seed EC. In addition, 140 other datasets from various other conditions or tissues were used in the global EC (Supplemental Data S11). All datasets were individually processed in R Bioconductor using a custom CDF file for Arabidopsis (Harb et al., 2010). The re-annotated CDF assigns probe-sets to specific genes and increases the accuracy in expression quantification. Using Robust Multi-array average algorithm (Irizarry et al., 2003), probe level expression values were background corrected, normalized and summarized into gene level expression values. Values from replicate arrays were then averaged and assembled in an integrated expression matrix of genes as rows and samples as columns, with each cell in the matrix representing log transformed expression value of genes in the corresponding samples. This procedure resulted in two expression matrices: a seed-specific expression matrix and a global expression matrix.

#### Coexpression network and clustering

Pearson’s correlation coefficients were calculated for each gene pair using expression values in both gene expression matrices separately. PCs were Fisher Z transformed and standardized to a N(0,1) distribution, where a Z-score of a gene-pair represents the number of standard deviations the observed coexpression score lies away from the mean of the dataset (Huttenhower et al., 2006). The following procedure was applied only to the seed network. Gene pairs with Z scores above 1.96 (PC 0.75) were retained and connected to create a coexpression network with 21,267 genes connected with approximately 7.6 million edges. SPICi was chosen as the clustering algorithm (Jiang and Singh, 2010), and used to cluster the network at a range of *T_d_* values ranging from 0.1 to 0.90, keeping a minimum cluster size of 3. Each *T_d_* value was evaluated on three criteria: i) total number of clusters yielded and the fraction of original genes retained in those clusters ii) average modularity following the (Newman and Girvan, 2004) algorithm and iii) functional coherence of clusters based on enrichment of GO BP term annotations (see next section). At *T_d_* 0.80, expression values of each gene within each of 1563 clusters were averaged across the same parts of the seed, as well as in different developmental stages, resulting in two expression profiles for each module. Expression values were scaled and plotted as heatmaps in R using the gplots package (https://CRAN.R-project.org/package=gplots).

#### Functional annotations of coexpression clusters

The TAIR gene association file was downloaded from the plant GSEA website (http://structuralbiology.cau.edu.cn/PlantGSEA/download.php) (Yi et al., 2013). The GO .gmt file was filtered to remove generic terms that annotate more than 500 genes, and the remaining list of terms in the BP category were used for testing overlaps with clusters. The KEGG .gmt file was similarly processed. The significance of overlap of a target gene set (e.g. a module) with a given BP term/KEGG pathways was calculated using a cumulative hypergeometric test. The *p*-values obtained were adjusted for false discovery rate and converted to *q*-values using the Benjamini-Hochberg method (Benjamini and Hochberg, 1995). Enrichment scores were reported as (−1) * log (*q*value).

#### Enrichment of CREs

We used a pattern-based method to search for CREs over-represented in the promoters of genes within the each module. First, all known plant motifs were identified from PLACE (Higo et al., 1999) and AGRIS databases (Palaniswamy et al., 2006). Subsequently, 1000-bp upstream promoter regions of all Arabidopsis genes were downloaded from TAIR and scanned for occurrence of these motifs using DNA-pattern matching tool (Medina-Rivera et al., 2015), yielding a list of 403 motifs present at least once in the promoters of ~17000 genes. A few of these motifs, perhaps involved in functions common to all promoters, are ubiquitously present in almost all genes. To detect a reliable presence-absence signal in the context of our analysis, we removed motifs that were found in more than 50% of all genes considered in the network. Thus, a list of 341 unique motifs were used for enrichment (overlap) analysis using a hypergeometric test as described above.

#### Module Regulatory Network analysis

A list of 1921 Arabidopsis TFs was curated from the Plant Transcription Factor Database, the AGRIS database and the Database of Arabidopsis Transcription Factors (Guo et al., 2005; Yilmaz et al., 2011; Jin et al., 2014) and 1819 were useable for this analysis. From the coexpression matrix of genes with normalized coexpression scores, we removed non-TF genes from the columns and retained a TF-gene coexpression matrix. Specific correlation scores were calculated for each TF and target gene following the CLR algorithm (Faith et al., 2007) and as described previously (Ambavaram et al., 2014). This produced in a matrix with 1819 columns (TFs) and 21,267 rows (genes) and each cell of the matrix populated with a *Z* score stating the likelihood of interaction between the corresponding TF and target gene. The Parametric Analysis of Geneset Enrichment (PAGE) (Kim and Volsky, 2005) algorithm was then used recursively on each column of the matrix (each TF) using seed coexpression modules as gene sets. Hence, the final TF-module association network was a matrix *E*, with matrix element *Eij* representing enrichment score between TF *i* and module j. To provide a normal distribution for association scoring, we used only those modules that had more than 10 genes, as suggested by the authors of the PAGE algorithm. FDR was adjusted using the Benjamini and Hochberg procedure (Benjamini and Hochberg, 1995). A global regulatory network was constructed the same way as the seed regulatory network, except that 140 non-tissue specific datasets were used.

#### Differential regulation analysis

Differential regulation was estimated by calculating the difference in normalized module association score in the two TF-module association matrices (1819 TFs in columns X 278 modules in rows), one of the global network and the other one of the seed network. Elements of resulting differential matrix were then subtracted from the mean of the matrix and divided by the standard deviation to get a standardized *Z* score. The sum of absolute *Z* scores of each TF across all modules was then computed as the weighted degree. TFs were sorted by the absolute weighted degree and ranked from 1 to *n* (*n*= number of TFs in the final network, here 1819). Lower rank values (close to 1) indicated TFs with largest changes in association with seed modules and vice versa.

#### Decile enrichment tests

The 63 genes in VGS was created by mining PubMed abstracts using specific search terms like “Arabidopsis thaliana seed embryo transcription factor” in the PubTator tool. To be selected for the VGS, we required the gene to be mentioned in these abstracts and present in our list of TFs. The final list was then manually checked for accuracy. The decile enrichment test was recently formulated for testing the significance of a genome-wide ranking of autism related genes (Krishnan et al., 2016). To perform this test here, the ranked list of TFs obtained from the differential regulation analysis was first divided into 10 equal bins (deciles). The overlap between the VGS and the first decile TFs (top 10%) was then computed and evaluated for statistical significance using a one-sided binomial test. The expected proportion was computed by counting the fraction of first decile TFs reported to not have a seed morphological phenotype (*n*=176; Supplemental Table S12). This list was created by searching the chloroplast phenomics database (Ajjawi et al., 2010). The experimentally validated target TFs of TFs in the VGS were identified from the Arabidopsis regulatory network in AGRIS (Yilmaz et al., 2011) and tested for enrichment in the first decile using the same approach as for the VGS.

#### Metabolite and seed storage compound analyses on dry seeds

*Arabidopsis thaliana* Col-0 wild type and homozygous Salk mutant plants were grown as described (Schneider et al., 2016). Once all plants were bolting, the youngest opened flower growing on the main stem was marked and, after 21 days when most of the siliques were dry, the four siliques above and below this marked silique were collected to retrieve sufficient amount of material for dry seed analyses. The marked silique was also included. Silique walls were removed from samples and 1.000 mg (± 5%) of dry seeds were subjected to biphasic extractions to separate polar metabolites, lipids, and insoluble proteins and analyzed as described (Schneider et al., 2016). Four plants of each genotype were analyzed. Briefly, lipids and proteins were analyzed by GC-flame ionization detection (FID) and a fluorescent hydrophobic protein assay as described (Collakova et al., 2013). Lipid- and oil-derived fatty acids present in the organic phase were analyzed as fatty acid methylesters on an Agilent 7890A series GC-FID (Agilent Technologies, Santa Clara, CA) equipped with a 30-m DB-23 column (0.25 mm x 0.25 μm, Agilent Technologies). Seed storage proteins present in the insoluble interphase (Schneider et al., 2016) were determined using a plate-reader-based fluorescent Marker Gene Hydrophobic Protein Analysis Kit (Marker Gene Technologies, Inc., Eugene, OR) and bovine serum albumin as a standard according to the manufacturer’s recommendations.

Polar metabolites and amino acids present in the aqueous phase were analyzed by Agilent GC-MS and Waters ultra-performance liquid chromatography (UPLC), respectively, as described (Schneider et al., 2016). Briefly, 5% of the aqueous phase was used for amino acid and organic amine derivatization with the Waters AccQ-Tag™ Ultra Kit. Amino acids were analyzed on an H-class Acquity UPLC equipped with a 10-cm Waters AccQ-Tag™ Ultra C18 (1.7 μm x 2.1 mm) column (Waters Corporation, Milford, MA) and coupled to a fluorescent detector. Waters 10.2-min method was used to analyze free amino acids as described (Collakova et al., 2013). The remaining aqueous phase was used to analyze other polar metabolites as trimethyl silyl derivatives on an Agilent 7890A series GC equipped with a DB-5MS-DG column (30 m length × 0.25 mm × 0.25 μm with a 10-m pre-column, Agilent Technologies) and analyzed on an Agilent 5975C series single quadrupole MS as described (Collakova et al., 2013). PCA on correlations was performed using JMP Pro 13 (SAS Institute Inc., Cary, NC).

Network data was parsed using the Sleipnir library of functional genomics (Huttenhower et al., 2008), Network Analysis Tools (NeAT) (Brohee et al., 2008) and scripts written in R and Perl.

### Author Contributions

C.G., A.K. and A.P. conceived the idea. C.G. designed the experiments, conducted statistical analysis and drafted the manuscript. A.K. provided software. A.S, C.D. and E.C. performed experimental validations and contributed text. C.G. created the webserver with contributions from P.W. A.P. designed the experiments, acquired funding and coordinated research. All authors read the manuscript and contributed in interpretation of results.

## Acknowledgements

The authors are grateful to all members of the Pereira and Collakova labs for insightful discussions on the biology of seed development during the development of this project. The authors would also like to thank NSF for providing funds to support this study (award number MCB 1716844).

## Supporting Information

### Supplemental Datasets

**Supplemental Data S1**: Clustering parition of genes in the seed coexpression network.

**Supplemental Data S2**: GO BP enrichment of coexpression modules.

**Supplemental Data S3**: CRE enrichment analysis of seed modules using DNA-pattern coupled with hypergeometric test.

**Supplemental Data S4**: Seed-specific correlation scores (Z-scores >= 4;) between TF and target genes estimated using the CLR algorithm.

**Supplemental Data S5**: Top 5 TF regulators of each module in SANe, with their absolute association score estimated by PAGE.

**Supplemental Data S6**: All the TFs in Arabidopsis ranked according to the sum of differences of module association scores between the SANe and the global network.

**Supplemental Data S7**: List of genes that were a part of the validated gene set (VGS) used for enrichment analysis of our first decile genes.

**Supplemental Table S8**: Module enrichment analysis of FUS3, LEC1 and LEC2 mutants (GSE61686).

**Supplemental Data S9**: Summary of mutant lines analyzed as part of SANe validation and their metabolic phenotypes.

**Supplemental Data S10**: Biomass and metabolite analyses.

### Supplemental Figures

**Supplemental Figure S1A and S1B**: Expression patterns of ATDOF5.8 and ATMYB62 by genevestigator analysis.

**Supplemental Figure S2**: Expression pattern of Module M0003

**Supplemental Figure S3**: Expression patterns of ATMYB62 by genevestigator

**Supplemental Figure S4A and S4B**: Expression patterns of AT2G21530 and WRI1 by genevestigator analysis.

**Supplemental Figure S5A and S5B**: Expression patterns of AT3G46770 and KAN3 by genevestigator analysis.

**Supplemental Figure 6**: Principal Component Analysis (PCA) on correlations of metabolite levels of select mutants

